# Metal-Organic Framework Encapsulated Whole-Cell Vaccines Enhance Humoral Immunity against Bacterial Infection

**DOI:** 10.1101/2020.06.14.148452

**Authors:** Michael A. Luzuriaga, Fabian C. Herbert, Olivia R. Brohlin, Jashkaran Gadhvi, Thomas Howlett, Arezoo Shahrivarkevishahi, Yalini H. Wijesundara, Sundharamani Venkitapathi, Kavya Veera, Ryanne Ehrman, Candace E. Benjamin, Sarah Popal, Michael D. Burton, Molly A. Ingersoll, Nicole J. De Nisco, Jeremiah J. Gassensmith

**Affiliations:** Department of Chemistry and Biochemistry; Department of Biological Sciences; School of Brain and Behavioral Science; Department of Biomedical Engineering at The University of Texas at Dallas. 800 West Campbel Rd. Richardson, TX 75080; Department of Immunology, Institut Pasteur, Paris, France

## Abstract

The increasing rate of resistance of bacterial infection against antibiotics requires next generation approaches to fight potential pandemic spread. The development of vaccines against pathogenic bacteria has been difficult owing, in part, to the genetic diversity of bacteria. Hence, there are many potential target antigens and little *a priori* knowledge of which antigen/s will elicit protective immunity. The painstaking process of selecting appropriate antigens could be avoided with whole-cell bacteria; however, whole-cell formulations typically fail to produce long-term and durable immune responses. These complications are one reason why no vaccine against any type of pathogenic *E. coli* has been successfully clinically translated. As a proof of principle, we demonstrate a method to enhance the immunogenicity of a model pathogenic *E. coli* strain by forming a slow releasing depot. The *E. coli* strain CFT073 was biomimetically mineralized within a metal-organic framework (MOF). This process encapsulates the bacteria within 30 minutes in water and at ambient temperatures. Vaccination with this new formulation substantially enhances antibody production and results in significantly enhanced survival in a mouse model of bacteremia compared to standard inactivated formulations.

## Introduction

Pandemic and epidemic spread of bacterial infection was common until the advent of antibiotic therapies. Indeed, some of the largest epidemic death tolls are attributed to bacteria (*e*.*g*., Black Death, Cocoliztli epidemic, Russian typhus epidemic, the Asiatic cholera pandemic). Combined, just these four bacterial pandemics/epidemics resulted in more than 200 million deaths.^1^ While antibiotics have largely ended pandemic spread, it is clear that the evolution of antibiotic resistance has already begun to complicate the treatment of common bacterial infections like urinary tract infection (UTI). The primary causative agent of UTI is uropathogenic *E. coli* (UPEC) with approximately 80% of all community-acquired UTIs caused by UPEC.^2-3^ At least half of all women will develop a UTI in their lifetime and more than 40% of these individuals will experience recurrent infection.^4-5^ While men are less likely to develop UTI, their prognosis is often worse than women because the infection is associated with greater morbidity and mortality and typically requires longer courses of antibiotics to resolve the infection.^6-9^ The incidence and recurrent nature of UTI takes on more urgency given that 10–25% of uncomplicated UTI patient isolates are resistant to trimethoprim/sulfamethoxazole, a front-line antibiotic for treatment of UTI.^10-11^ When antibiotic treatment fails, bacteria persist in the bladder, increasing the cost of care and morbidity to the patient.^12^ Left untreated or undertreated, lower UTI can ascend to the kidneys and cross into the bloodstream, progressing to the more severe diseases of pyelonephritis and urosepsis, which have a global mortality rate of 40%.^13-14^ Given the rise in antimicrobial resistance among UPEC strains and its potential lethality, efforts to develop non-antibiotic based therapies for UTI have increasingly focused on the development of prophylactic and therapeutic vaccines against UPEC. However, vaccine development against bacterial infection is challenging. In fact, no effective vaccines against any type of pathogenic *E. coli* have been clinically successful thus far. Therefore, new strategies to improve the efficacy and immunogenicity of bacterial vaccines must be developed.

Commonly, vaccines that target bacteria are created in three different ways. The first includes live-attenuated strains that are less pathogenic but capable of mutation and inducing strong immune reactions themselves.^15-16^ A second method is to use specific surface antigens found on bacteria in a subunit approach. However, subunit vaccines against *E. coli* strains tend to elicit poor T cell responses^17-18^ and have yet to translate into effective vaccines against UTI in humans. Further, a weak T-cell response will hinder antibody production and class switching, a significant problem when a humoral response is necessary to fight UTI.^19-21^ One reason for this may be that because the surface of *E. coli* is compositionally mosaic^22^ and antigens present within the outer membrane differ greatly between strains. A recent sequencing approach identified 230 potential protein surface antigens, of which only nine were ultimately protective in mice.^23-24^ A third approach is to create vaccines based on whole or lysed fractions of inactivated (dead) bacteria. This whole cell approach has produced more promising results in humans compared to purified subunit vaccines.^25-26^ Whole cell vaccines contain both conserved and strain-specific proteins and, as such, should provide the most comprehensive protection. However, whole-cell vaccines often fail to be protective in the long-term. ^27-31^ The primary hurdle for the development of inactivated whole cell bacterial vaccines is the weak immune response they elicit because of their short half-life within the body, poor uptake by antigen presenting cells, and/or surface antigen degradation by harsh fixation methods.^32-34^

Methods to slow the release of antigens via an injected depot have profoundly positive effects on development of long-term antibody-based immunity by promoting germinal center (GC) formation in the draining lymph nodes.^*35-37*^ GCs are the primary site of B cell maturation into antibody-producing plasma B cells. Whereas thermoplastics,^38-43^ liposomes,^44-47^ micelles,^48-49^ porous silicon,^50-53^ and polypeptides,^54-55^ have shown promise for small protein release and antigen delivery,^56-60^ it is not clear how these technologies will translate to whole-cell formulations. Methods to fully encapsulate “large cargo” such as whole bacterial cells are less explored. To address this challenge, we developed biomimetic encapsulation^61-62^ methods that fully encase individual bacteria inside a crystalline polymeric matrix called <u>Z</u>eolitic Imidazolate Frameworks (ZIFs — **Scheme 1**).^63^ We and others showed that mixing an array of biological molecules, including fluorescent proteins,^64^ insulin,^65^ proteins,^66-67^ IgG,^68^ viral nanoparticles,^37, 69-70^ RNA, yeast,^71-72^ proteoliposomes,^73^ and Gram-positive and - negative bacteria^74^ within the ZIFs framework, incarcerates the target in a biodegradable matrix. ZIFs are a subclass of a broader family of solid-state single crystal metal organic frameworks (MOFs) characterized by interconnected organic ligands linked by metal nodes extending infinitely in all directions (**Scheme 1A and 1B**). We found that ZIF-8, a metal-organic framework composed of zinc ions interlinking 2-methylimidazole (HMIM) molecules in a repeating framework (**Scheme 1B**) self-assembles on protein surfaces through weak interactions between Zn^2+^ and peptide backbones.^70^ The resulting ZIF-8 shell (**Scheme 1C**) has exceptional thermodynamic stability against high temperature, moisture, and organic solvents, but is kinetically labile in the presence of metal chelators such as EDTA and inorganic phosphates. These characteristics make ZIF-8 an ideal material for reversible immobilization.^71, 75-76^ Here we report that, ZIF encapsulation preserves natively folded bacterial proteins better than traditional methods of preparing inactivated bacterial vaccines. Further, post-injection, ZIF-coated whole cell UPEC dissipate slowly at the injection site, induce a stronger humoral response compared to the controls, and provide superior protection against fatal septicemic UPEC infection in mice as compared to existing literature approaches.^37^

**Scheme 1:**
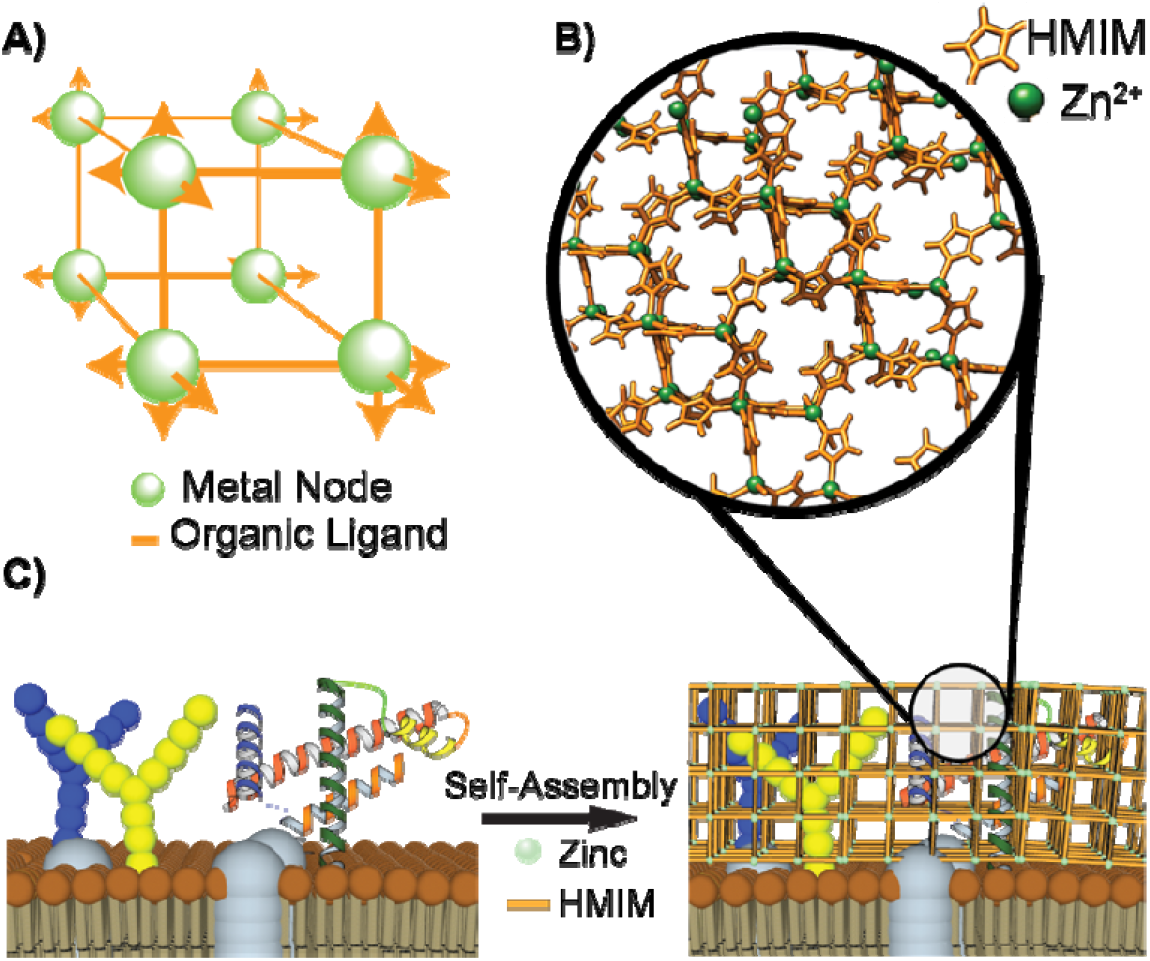
A) Simplified schematic of a metal organic framework composed of metal nodes and organic ligands that can expand infinitely in all directions. B) The crystalline super structure of ZIF-8 is illustrated with the orange HMIM ligands connecting the green zinc ions. C) Conceptualization of the synthetic process shows that the surface of an *E. coli* external membrane can initiate ZIF-8 growth on and around membrane-bound biomacromolecules following incubation with Zn^2+^ and 2-methylimidazole (HMIM).

## Results and Discussion

### Uropathogenic E. coli support growth of ZIF

For this study, we used CFT073, a well-studied urosepsis strain of uropathogenic *Escherichia coli* (UPEC) frequently employed in the development of new vaccines, to allow us to benchmark our work against the literature.^77^ Gram-negative *E. coli* have an outer membrane containing embedded polysaccharides and proteins,^20, 23, 78^ which support ZIF-8 growth. We resuspended CFT073 in a saline solution containing 2-methylimidazole (HMIM) and zinc acetate and after washing to remove excess zinc and HMIM, analyzed the resulting encapsulated bacteria, called CFT@ZIF, by scanning electron microscopy (SEM), powder X-ray diffraction (PXRD), and energy dispersive X-ray (EDX) spectroscopy. ZIF-8 growth occurred exclusively on the surface of the bacteria (**Figure 1A**). The PXRD of CFT@ZIF confirmed the shell was crystalline ZIF-8 (**Figure 1B**) and EDX confirmed the presence of zinc and phosphorus, indicating both ZIF-8 and bacteria, respectively, were in the preparation (**Figure 1C and D**).

**Figure 1.**
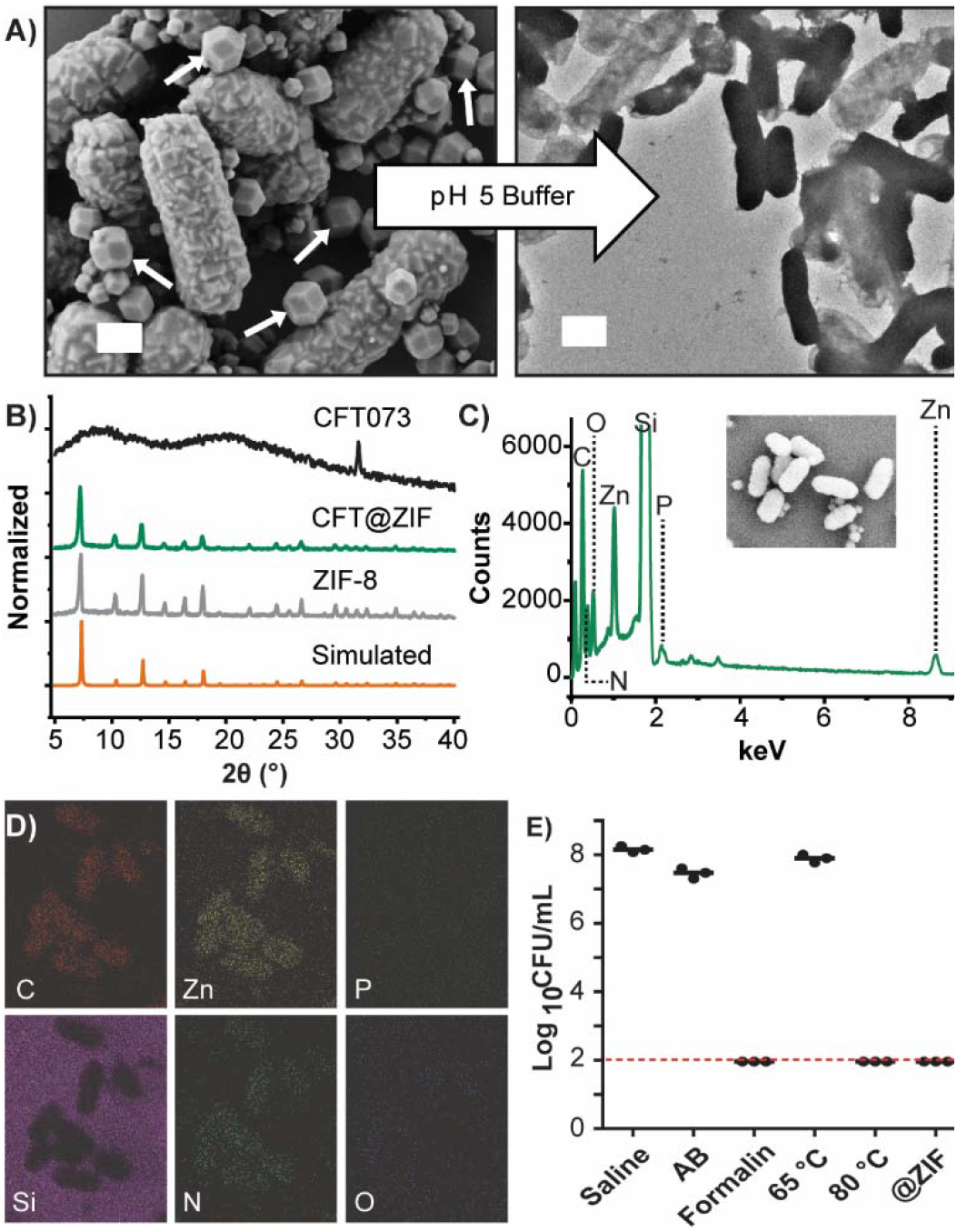
Physical characterization of CFT@ZIF: A) SEM micrograph of CFT@ZIF (left). The ZIF shell is removed in sodium acetate buffer (AB) at pH 5 to reveal intact bacteria, as seen in the transmission electron micrograph (right). White scale bars are 1 µm and white arrows are free ZIF crystals. B) PXRD of CFT@ZIF compared to pristine/empty ZIF-8, showing the measured data matches simulated PXRD spectra of pristine ZIF-8—CFT073 was ran as an additional control. C) Element distribution of CFT@ZIF by EDX. The graph shows the presence of carbon, oxygen, nitrogen, zinc, and phosphorus. D) Image maps show zinc, oxygen, nitrogen, zinc, and phosphorus signals come from CFT@ZIF. E) Bacteria growth assay shows CFT@ZIF is not viable after exfoliation, like formalin fixation or heat treatment, and can be used as a method to inactivate bacteria. The dashed line indicates the detection limit of 100 CFU/mL. It should be noted that bacteria incubated in acetate buffer (AB) at pH 5 was used as control to show that the acidic buffer used to deshell ZIF causes no harm to growth.

An initial challenge was the formation of free or empty ZIF-8 crystals. To find conditions that reduced the number of empty ZIF-8 particles (**Figure 1A**, white arrows), we tested a time course of incubation and a range of salinity in the encapsulation solution. We observed that an encapsulation time of 20 minutes and a final concentration of 100 mM NaCl yielded crystal growth primarily on the bacteria (**Fig. S1A-H**). Using these parameters to generate CFT@ZIF, we then tested whether we could remove the shells from the encapsulated bacteria, in a process called exfoliation. We observed by transmission electron microscopy that the bacteria remained intact after dissolving the ZIF-8 shell in mildly acidic (pH 5) sodium acetate buffer and no obvious debris from lysed bacteria were seen (**Figure 1A**). Next, we tested the stability of CFT@ZIF when lyophilized or kept in 0.9% saline solution. Both formulations were monitored over the course of 28 days by taking SEM images, which showed the composites had no obvious changes (**Fig. S2A**,**B**).

A challenge in whole-cell vaccine formulation is inactivating the bacteria with minimal damage to the surface epitopes, including delicate membrane-bound proteins and complex oligosaccharides. Biomimetic mineralization with ZIF is extremely gentle and we have previously shown that membrane bound proteins are not only preserved but protected by ZIF encapsulation in lipid nanoparticles.^73^ To show that our ZIF encapsulation approach preserves natively folded proteins better than traditional methods of inactivation, we conducted an agglutination assay using yeast. The fimbriae on the surface of CFT073 contain glycoproteins that are highly specific binders of sugar. When these proteins are damaged, they cannot bind sugar. Yeast, which have cellular surfaces rich in mannoprotein will crosslink bacteria forming aggregate clumps that can be directly imaged via microscopy as shown in the illustration in **Fig. S2C**. In our experiment, only untreated CFT073 and CFT@ZIF retained binding to the yeast, whereas CFT-Fixed and CFT-heat treated did not show any binding at the same concentration (**Fig. S2D**). These results support the that the growth of ZIF does not damage surface bound molecules. In addition, CFT@ZIF was left undisturbed in saline solution for 7 days and the agglutination was tested at day 0 and 7, which showed after 7 days it still retained binding to the yeast (**Fig. S3**). We next tested whether ZIF encapsulation inactivates CFT073. Following formation of the ZIF shell, immobilized CFT073 was incubated for 30 min at room temperature and exfoliated with 500 mM sodium acetate buffer at pH 5. Unencapsulated CFT073 was inactivated using formalin or heating as described for other whole-cell bacterial vaccines,^79-81^ and bacterial viability was measured for each condition by colony forming unit (CFU) assay, and compared to untreated CFT073 incubated in 0.9% saline. Bacterial growth was observed for untreated CFT073 incubated in saline or in the sodium acetate buffer but not for bacteria that were formalin-fixed, heat-inactivated, or, importantly, exfoliated from the ZIF shell (**Figure 1E**). These results support that the ZIF-shell growth process itself inactivates bacteria, which is in line with previous observations that overexposure to zinc causes bacterial cell death.^82-84^

### Vaccination with CFT@ZIF induces a robust IgG response

Having established that CFT073 was encapsulated, intact, and inactivated, we turned to vaccination studies. A humoral response depends on the activation of antigen-specific B cells to differentiate into antibody-producing plasma B cells.^85^ We hypothesized that the ZIF shell would induce higher antibody titers because it not only avoids chemical alterations and thermal denaturation of surface proteins but also creates a slow releasing depot. We benchmarked our CFT@ZIF vaccine formulation against known methods to inactivate bacteria for whole-cell vaccine use: formalin fixation and heat treatment.^80, 86-87^ Formalin fixation induces chemical crosslinking of proteins to preserve their tertiary structure, resulting in bacterial cell death, whereas heat treatment kills bacteria through thermal stress.^88^ Here, formalin-fixed CFT073 are referred to as CFT-fixed and heat treated CFT073 are denoted as CFT-Heat. We first compared the time-resolved fluorescence of smURFP-expressing CFT-Fixed against CFT@ZIF. The near-infrared fluorescent protein smURFP was chosen for its durability and brightness and has been used to image nanomaterials in tissues as deep as the kidney *in vivo*. During heating, the smURFP expressed within CFT lost fluorescence, so CFT-heat was not used in the following experiments. Mice were injected with either saline, or the smURFP-expressing CFT-Fixed, or CFT@ZIF preparations subcutaneously and monitored over a period of 12 days until fluorescence levels returned to baseline (**Figure 2A**). Images collected by monitoring smURFP fluorescence during this period show that intact CFT@ZIF (t_1/2_ = 106.4 h) remained at the injection site 4 days longer than CFT-Fixed (t_1/2_ = 11.1 h, **Figure 2B**).

**Figure 2.**
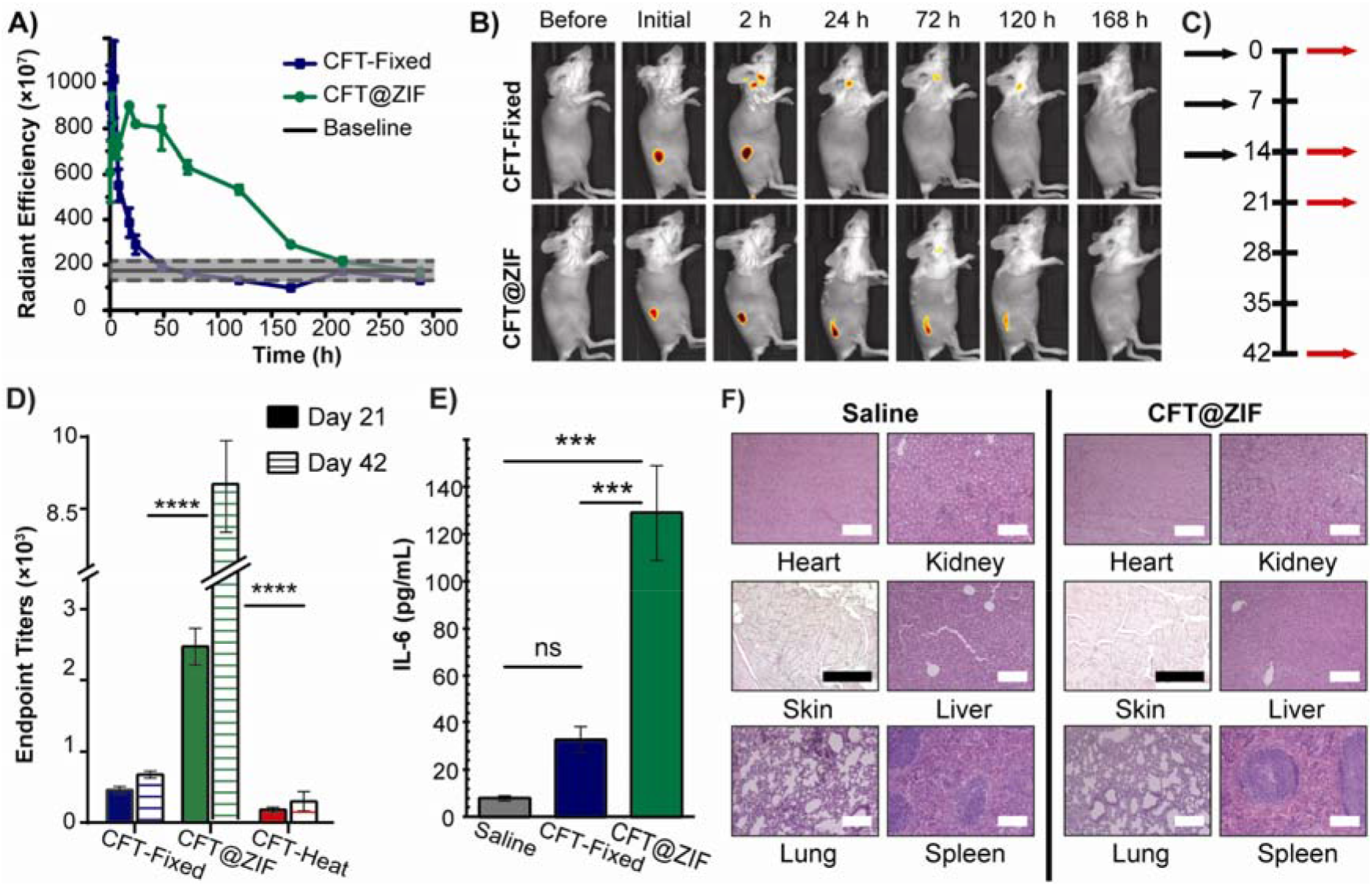
*In vivo* release: BALB/c Mice (n=4) were injected with smURFP-expressing CFT073 that was inactivated with formalin (CFT-Fixed) or by encapsulation (CFT@ZIF). A) The graph shows fluorescence change at the site of injection over time. The baseline is the average fluorescence of four mice before and after injecting with saline. The dashed line represents the error of the baseline. B) Representative images of mice prior to injection with CFT-Fixed or CFT@ZIF and after injection were monitored over a course of 12 days. After injection, images were taken at 15, 30, 60, 120, 240, and 480 min, then every 12 h for 12 days. C) Vaccination schedule for mice injected with either saline, CFT-Fixed, CFT@ZIF, or CFT-Heat. Black arrows are vaccinations and red arrows are blood draws. D) At day 21 and 42, blood was drawn and measured by ELISA to determine the antibody production. E) IL-6 in serum at day 14. F) H&E staining of organs at day 42. Error bars represent the mean±SD. Statistical significance was calculated using an ordinary one-way ANOVA with Tukey’s multiple comparison post-test (*p < 0.05, **p < 0.01, ***p < 0.0005, ****p < 0.0001). White scale bars are 100 µm and black scale bars are 200 µm.

We next measured antibody titers in mice injected with saline, CFT-Fixed, CFT-Heat, or CFT@ZIF according to the vaccination and blood sampling schedule in **Figure 2C**. Serum from days 21 and 42 post-injection was serially diluted and the amount of anti-CFT073 IgG produced was determined by enzyme-linked immunosorbent assay (ELISA). Mice immunized with CFT@ZIF produced the highest levels of anti-CFT073 IgG at day 21 and 42 compared to all other groups (**Figure 2D**). As expected, the thermally-inactivated CFT-Heat formulation induced the lowest antibody production, in line with the observation that high temperatures lead to thermal denaturation of proteinaceous and sugar-based epitopes.^89^ Following productive interaction with Th cells, activated B cells undergo antibody class switching and change from producing IgM to IgG. IgG antibody responses can be further divided into subclasses, depending on the types of antigens they sense. We measured the different IgG subclasses by running the serum from day 21 on a multiplex assay, which showed an increase of IgG1, IgG2a, and IgG2b for CFT@ZIF compared to CFT-Fixed (**Fig. S4**). B cells tend to switch to IgG2a when exposed to the inflammatory Th1 cytokine IFN-γ or other proinflammatory agents such as LPS, whereas IgG2b and IgG1 are produced in the presence of the Th2 cytokine IL-4 and TGF-β. Thus, our results suggest a stronger T cell presence for CFT@ZIF compared to CFT-Fixed. As IL-6 promotes production of immunoglobulins by plasma cells,^90^ we diluted serum from day 14 and measured IL-6 production by ELISA. IL-6 levels were highest in CFT@ZIF, which correlated with enhanced production of antibodies in mice injected with this formulation (**Figure 2E**). The increase in CFT073-specific antibodies from mice injected with CFT@ZIF combined with the prolonged residency of CFT@ZIF at the injection site and the retained binding in the agglutination assay, supports the hypothesis that ZIF encapsulation helps prevent protein denaturation on the surface and provides a depot effect by prolonging the presence of CFT@ZIF in the tissue compared to unencapsulated bacteria.

ZIF-8 is generally non-toxic^91-92^ and, throughout our experiments, mice monitored daily showed no signs of pain or changes in behavior. At day 42, the mice were euthanized, and the spleen, liver, kidney, injection site tissue, lung, and heart were collected, fixed in formaldehyde, and stained with hematoxylin and eosin (H&E) for pathological analysis. Organs displayed no signs of toxicity and H&E-stained organ sections showed no abnormal lesions, aggregation, or change in tissue compared to mice injected with saline (**Figure 2F**) indicating the CFT@ZIF vaccine did not induce acute toxicity.

Cellular responses in the spleen and draining lymph node were characterized by flow cytometry on day 21 following the vaccination schedule illustrated in **Figure 2C**. Notably, T-cell populations (CD3+, CD4+, and CD8+) were significantly elevated in the spleen of CFT@ZIF-treated mice compared to animals receiving the CFT-fixed formulation (**Figure 3A**) whereas T-cell responses against both formulations were comparable in the draining lymph node (**Figure 3B**). These data suggest that the CFT@ZIF formulation promotes a systemic immune response—as indicated by the more robust response in the spleen. Notably, B-cell numbers (CD19+) were significantly elevated in both the spleen and draining lymph node after CFT@ZIF vaccination, which was expected given the strong IgG response this formulation produced. We hypothesize that these results are a direct result of the slow-release depot effect, which we assess in the following section.

**Figure 3.**
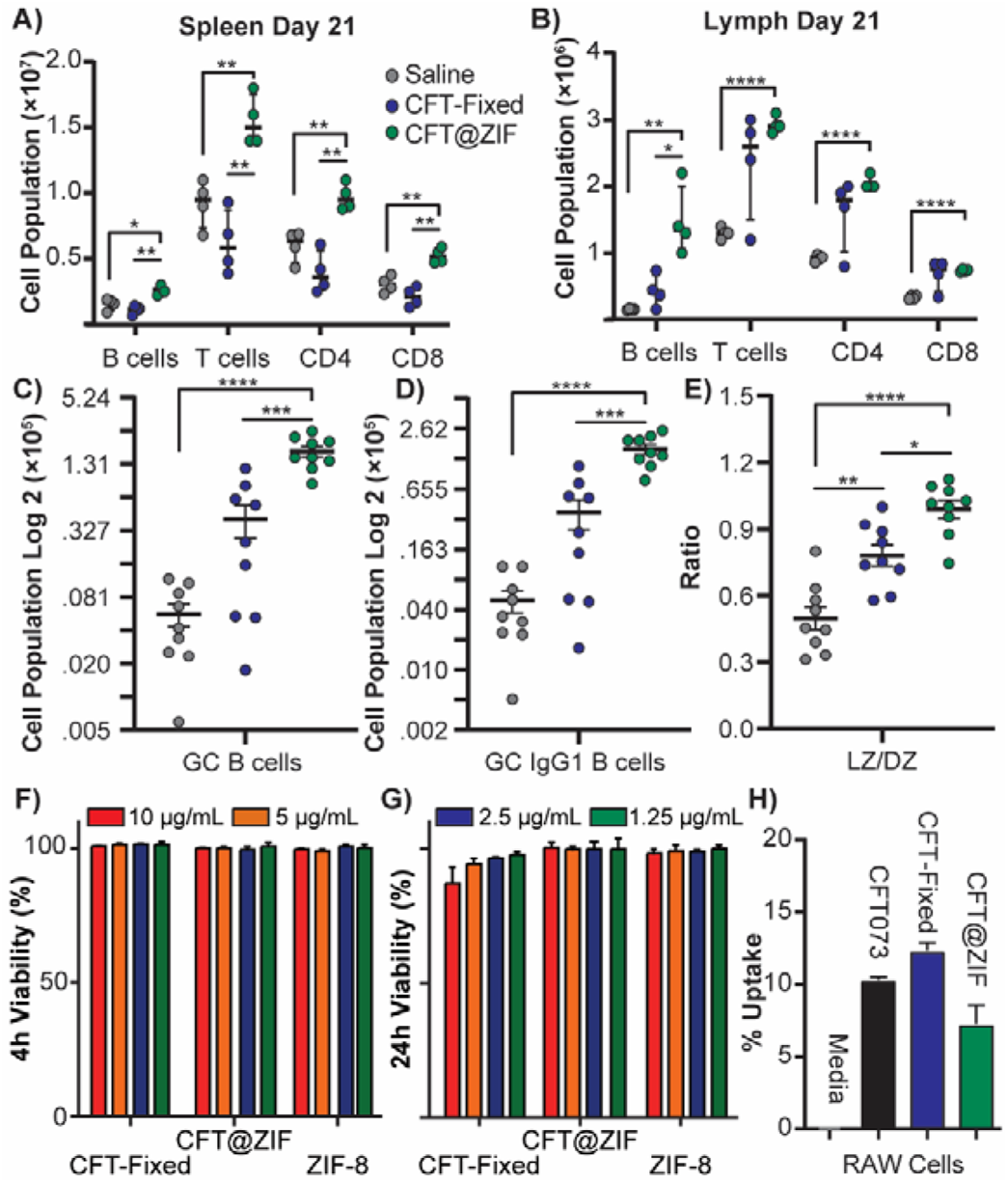
Cellular response to different formulations: immune cell population of A) the spleen and B) draining lymph node (N = 4) at day 21 after following the vaccination schedule in Figure 2C. A second (N = 3) and third cohort of mice (N = 6) were vaccinated following schedule in **Figure 2C**, to determine C) the germinal center B cells, D) the IgG1^+^ B cells within the germinal B cells population, and E) the ratio of LZ to DZ in the germinal center of the draining lymph node. The viability of CFT-Fixed, CFT@ZIF, and ZIF was tested at different concentrations on T24 urinary bladder carcinoma cell line for F) 4 h and G) 24 h. H) Bar graphs showing true cellular uptake of different CFT073 formulations. Error bars represent the mean±SD. Statistical significance was calculated using one-way ANOVA with Tukey’s multiple comparison post-test (*p < 0.05, **p < 0.01, ***p < 0.0005, ****p < 0.0001).

Antigen persistence via the depot effect is correlated with induction of a strong adaptive immune response.^35, 93^ In a traditional bolus immunization, the half-life of the injected antigen is thought to be shorter than the required development time of germinal centers (GCs) in peripheral lymph tissue.^35^ GCs are transient anatomical structures that form in lymphoid organs when early antigen-stimulated B-cells migrate and diversify into affinity B-cells and durable memory B-cells. The formation of GCs takes approximately a week and it is hypothesized a slow yet persistent release of antigens better matches the kinetics of GC B-cell development and ultimately the selection process for B-cells that produce high-affinity antibodies.^94^ In other words, by continuously supplying a source of antigens, GC B-cells have a longer opportunity to evolve the high affinity antibodies necessary to neutralize infections and if our depot mechanism is having an effect, we should see improved GC development. GC development can be assessed by measuring the amount of GC B-cells (GCBCs—CD19+, CD95+, and GL7+) in lymph tissue. To perform this analysis, a second cohort of mice were vaccinated following the schedule in **Figure 2C** and on day 21, the draining lymph nodes were removed. We saw a larger population of GCBCs in the draining lymph nodes of mice vaccinated with CFT@ZIF compared to fixed CFT (**Figure 3C**) and a higher frequency of class switched GCBCs (IgG1^+^) (**Figure 3D**), which is strongly correlated to a protective antibody response. GCs develop into two different microenvironments—a dark zone (DZ), where GCBCs undergo somatic hypermutation and create receptors of varying affinity against an antigen, and a light zone (LZ), where only antigen-specific high-affinity receptor B-cells are selectively differentiated into memory or plasma B cells.^95^ Consequently, LZ growth indicates an increase in affinity selection and production of B-cells that produce high affinity antibodies. When we analyzed the draining lymph nodes by flow cytometry, we found a shift in the light zone to dark zone (LZ/DZ) ratio for mice vaccinated with CFT@ZIF (**Figure 3E**) compared to the formalin inactivated formulations. Taken together, we postulate that a likely reason we see such a strong B cell response from CFT@ZIF is from the antigen depot effect.

A second mechanistic explanation for the higher antibody titers may be increased uptake by professional phagocytes. Prior literature has suggested that ZIF-8 improves cellular uptake.^96-99^ ZIF-8 dissolves in the acidic compartments of phagosomes, endosomes, and lysosomes, which would free surface epitopes in a manner similar to acidic exfoliation buffer. First, we tested the toxicity of CFT@ZIF by conducting an LDH assay at 4 h and 24 h at 1.25, 2.5, 5, 10 µg/mL with T24 human urinary bladder carcinoma cell line and RAW 264.7 macrophages (**Figure 3F**,**G** and **Fig. S5A**,**B**), which showed no toxicity at all concentration tested and is in line with other biocompatibility studies using ZIF^91-92, 96^—CFT-Fixed and ZIF were used as controls. Next, we performed ZIF-8 shell growth on smURFP-expressing CFT073. CFT073, CFT-Fixed, and CFT@ZIF were incubated for four hours with RAW 264.7 macrophages to investigate if our ZIF encapsulation improved uptake. Confocal microscopy images showed that CFT@ZIF was taken up more readily by the RAW 264.7 macrophages (**Fig. S5C**) and flow cytometry analyses found a four-fold increase in uptake of CFT@ZIF over untreated CFT073 and CFT-fixed (**Fig. S5D**). While this initially appeared to validate prior literature observation, upon closer inspection, we found a significant amount of the CFT@ZIF had adhered to the surface of the macrophages. To remove this surface-attached ZIF-encased bacteria, we included three rapid washes with a low pH buffer—a method developed originally to strip IgE antibodies from cell surfaces.^100-102^ Another LDH assay was conducted to ensure that the low pH wash would not be toxic to the RAW 264.7 macrophages, which again showed the treatment did not kill the cells (**Fig. S5E**). The low pH wash completely dislodged the ZIF bound to the macrophage surface as confocal micrographs show no CFT@ZIF on the surface of the acid-washed macrophages (**Fig. S5F**). Following our acid washing step, flow cytometry revealed that the uptake of CFT@ZIF was lower than the other washed inactivated samples (**Figure 3H**), further suggesting that a depot effect is likely the principal driver of the increased immune activation.

### Vaccination with CFT@ZIF protects mice from lethal sepsis challenge

Activation of a cell-mediated adaptive immune response is a vital aspect of vaccine development to promote long-term memory.^103-104^ To determine whether CFT@ZIF promotes a cell-mediated adaptive immune response, we measured TNF-α, IL-2 and IFNγ serum levels. TNF-α is an immunostimulatory cytokine mainly produced by macrophages and T_h_1 (Type 1 T helper) CD4^+^ T cells while IL-2 is a cytokine produced mainly by CD4+ T cells and activated CD8+ T cells to promote T cell survival and T cell differentiation. IFN-γ is a cytokine produced mainly by T_h_1 T cells and dendritic cells promoting a cell-mediated response by stimulating cytotoxic T lymphocytes.^105-106^ In addition, we measured IL-4, a cytokine produced in part by T_h_2 (Type 2 T helper) CD4+ T cells, which induces an antibody-mediated response and IL-17, a cytokine that plays a protective role against *E. coli* infection.^107-109^ We employed the vaccination schedule depicted in **Figure 2C**, and on day 42, re-stimulated splenocytes with CFT073 for each group. We measured TNF-α, IL-2, IFN-γ, IL-17, and IL-4 cytokine (**Figure 4A-E**) levels by ELISA and observed that mice immunized with CFT@ZIF had higher levels of all cytokines compared to CFT-Fixed treated mice following re-stimulation. In particular, mean titers of IL-2 and IL-17 were more than 2-fold higher in splenocytes derived from CFT@ZIF compared to CFT-Fixed vaccinated mice (**Figure 4B,D)**. Titers of TNF-α, IFN-γ, and IL-4 were 1.4-fold, 2.0-fold and 2.0-fold higher, respectively, in splenocytes derived from CFT@ZIF vaccinated mice compared to CFT-Fixed vaccinated mice (**Figure 4A, C**,**E**). Taken together, our data demonstrate that CFT@ZIF induces a more robust cytokine response that is indicative of enhanced T cell activation.

**Figure 4.**
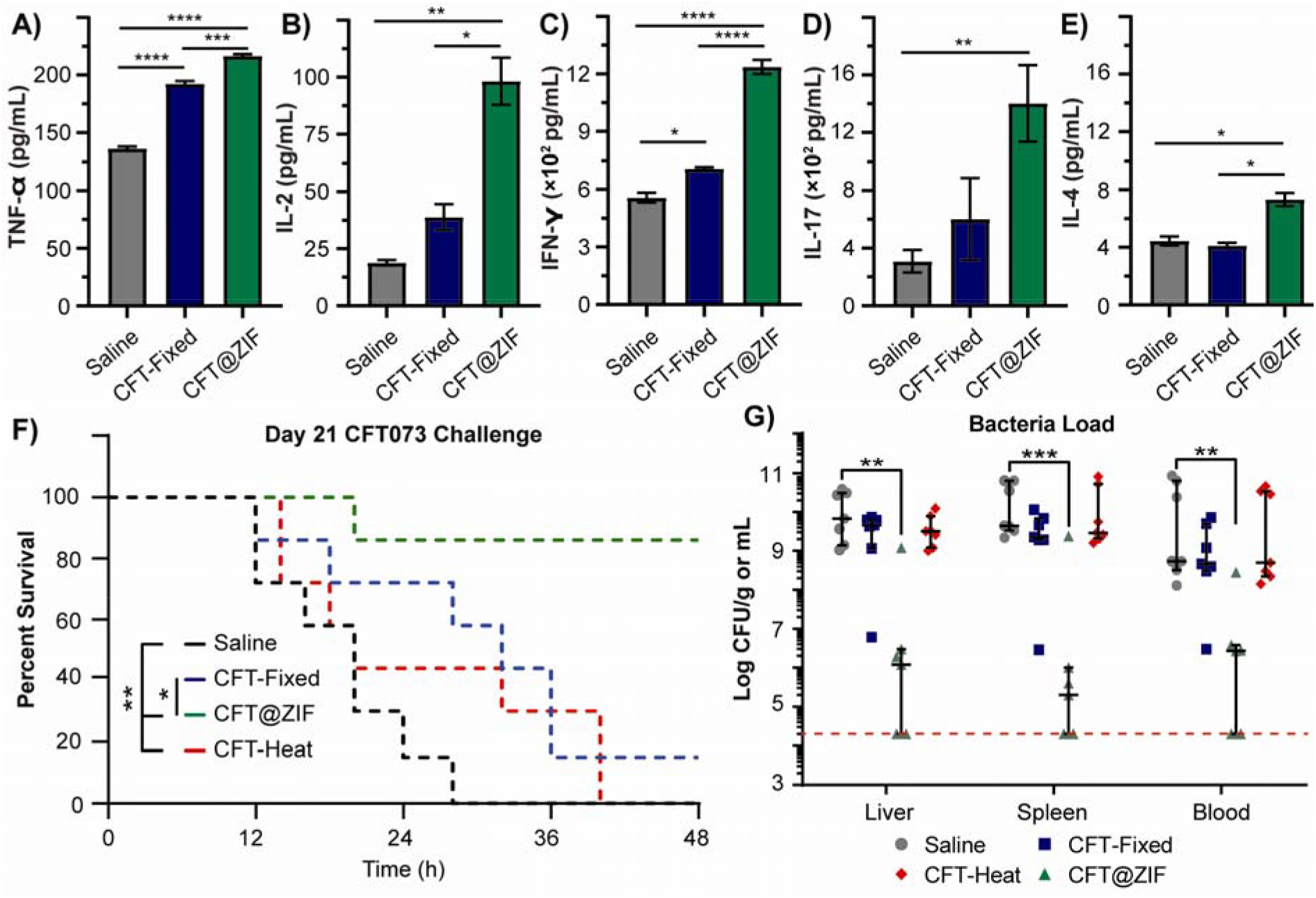
Survival Study: Mice (n=4) were injected with CFT073 that was inactivated with formalin (CFT-Fixed) or by encapsulation (CFT@ZIF). At Day 42, splenocytes were collected from immunized mice and incubated with 10 µg/mL of CFT073 for 48 h. After 48 h, the supernatant was tested for A) TNF-α, B) IL-2, C) IFN-γ, D) IL-17, and E) IL-4. Two separate cohorts of mice (n=3 and n=4) following the vaccination schedule in **Figure 2C** were injected interperitoneally with a lethal dose of CFT073 at day 21 and monitored for 48 h. F) Survival for each group over the course of 48 h. G) Bacterial loads in the liver, spleen, and blood at the endpoint of the survival study. Error bars represent the mean±SD. Statistical significance was calculated using an ordinary one-way ANOVA with Tukey’s multiple comparison post-test (*p < 0.05, **p < 0.01, ***p < 0.0005, ****p < 0.0001).

Mice vaccinated with CFT@ZIF had increased antibody production and increased expression of key cytokines at day 42 compared to other CFT formulations (**Figure 4A-E**). To determine whether these responses would be protective, we used a previously described urosepsis model.^12, 110^ Two survival studies performed with separate preparations of vaccine composites were conducted. We vaccinated as shown in **Figure 2C** with either saline, CFT-Fixed, CFT-Heat, or CFT@ZIF. On day 21, we administered a lethal dose^110^ of CFT073 intraperitoneally and mice were monitored over the course of 48 hours and euthanized when they became moribund, defined by lack of movement for over 15 minutes, shaking in place, and a decrease of body temperature or the 48-hour endpoint was reached. The Kaplan Meier analysis reported in **Figure 4F** shows that CFT@ZIF-vaccinated mice had improved survival compared to all other groups. The median survival time for sham treated (saline) controls was 16 hours compared to 20 hours for mice vaccinated with CFT-Heat and 32 hours for mice vaccinated with CFT-fixed. Strikingly, 85.7% (6/7) of the CFT@ZIF-vaccinated mice survived until the 48 hour endpoint with no visible signs of disease. After euthanasia, blood, spleens, and livers were collected, homogenized, and CFUs were enumerated (**Figure 4G**). Average bacterial load in the blood (2.7×10^6^ CFUs), spleens (1.2×10^6^ CFUs), and livers (2.0×10^5^ CFUs) of CFT@ZIF-vaccinated mice were three-to-four orders of magnitude lower (∼10^9^–10^8^ CFUs) than all the other groups with three CFT@ZIF animals having no detectable bacterial burden in the blood, spleen, or liver. It is noteworthy that this reduction in bacterial load and survival exceeds the efficacy of published subunit vaccine strategies for CFT073 sepsis.^12^ We hypothesize that the lower bacterial load and greater survival of the CFT073@ZIF-vaccinated mice may be attributed to the 5-fold higher anti-CFT073 IgG titers that CFT@ZIF promotes over the more traditional CFT-Fixed formulation (**Figure 2D**).

## Conclusion

In this work, we demonstrated that ZIF-8 effectively inactivated a urosepsis strain of UPEC, creating a persistent vaccine depot that considerably outperformed current whole cell inactivation methods in survival following lethal challenge in a sepsis model, B-cell, and T-cell activation. This method of inactivation avoids the use of toxic formaldehyde, is faster, and yields consistent superior results. Only CFT@ZIF-vaccinated mice survived a lethal dose of CFT073, which we hypothesize may be due to the five-fold increase in serum levels of IgG that arises from our depot approach, as evidenced by greater GC formation in draining lymph tissue. Finally, the biomimetic growth strategy that we employed here is likely generalizable across different organisms, presenting an opportunity for a generalizable approach toward whole-cell bacterial vaccine formulation. Formulation of the inactivated bacteria does not use dangerous or toxic compounds, involves the mixture of only three components, and the total reaction time is under 30 minutes. We and others have shown, moiety encapsulation with ZIF is a straightforward method that can easily be scaled up. Further, increasing reaction volumes does not alter the morphology of the as obtained materials as shown by PXRD (Powder X Ray diffraction) and SEM.^74, 111^ Our ecologically friendly and low-cost approach has shown that it can produce protective effects that exceed published subunit strategies making it an ideal platform for rapid vaccine production from patient-derived bacterial strains. This opens potential opportunities to produce patient-specific prophylactic and therapeutic vaccine formulations for those suffering from recurrent UTI or at increased risk for complicated UTI in a personalized medicine approach.

## Methods

### Materials

Acetic acid, acetic anhydride, anti-mouse IgG (whole molecule)-alkaline phosphatase produced in goat, arabinose, bovine serum albumin, diethanolamine, magnesium chloride, β-mercatoethanol, methanol, 2-methylimidazole, paraformaldehyde, p-nitro phenyl phosphate, potassium hydroxide, potassium phosphate dibasic, potassium phosphate monobasic, poly(vinylpyrrolidone) 40k (PVP 40k), 2-propanol, sodium azide, sodium bicarbonate, sodium carbonate, sodium chloride, sodium hydroxide, sodium phosphate dibasic, sodium phosphate monobasic, Tween-20, and zinc acetate dihydrate were purchased from Sigma-Aldrich (St. Louis, MO, USA), Thermo Fisher Scientific (Waltham,

MA, USA), Chem-Impex Int’l (Wood Dale, IL, USA), or VWR (Radnor, PA, USA), and used without further modification. Anti-rat CD3-APC, anti-mouse CD4-PE/Cy7, anti-rat CD8a-APC/Cy7, anti-mouse CD44-FITC, anti-rat CD62L-BV605, ELISA MAX™ Standard Set Mouse IL-6, TNF-α, and IFN-γ were from Biolegend. LDH-Cytox™ Assay Kit was purchased from Biolegend (Cat. No. 426401). The LEGENDplex™ Mouse Immunoglobulin Isotyping Panel (6-plex) was bought from BioLegend. Ultrapure water was obtained from an ELGA PURELAB flex 2 system with resistivity measured to at least 18.2 MΩ -cm.

### Bacterial Studies

#### Bacterial strains and growth conditions

CFT073 was obtained from ATCC and grown in Luria Bertani (LB) medium at 37°C for all experiments. pLenti-smURFP was a gift from Erik Rodriguez & Roger Tsien and was transformed into CFT073 electrocompetent cells as described: 50 µL of electrocompetent CFT073 were thawed on ice, then mixed with 100 ng of pLenti-smURFP and incubated on ice for 20 minutes. The mixture was transferred to an electroporation cuvette and electroporated at 1.8 kVs. 500 µL of SOB media was immediately added and the cells recovered for 60 minutes at 37°C. Transformants were selected on LB agar plates supplemented with 100 µg/mL ampicillin. For encapsulation experiments, CFT073-smURFP was grown to an OD_600_ of 0.9-1 and 10 mg/mL arabinose was added to induce smURFP expression. After 4 hours, cells were harvested by centrifugation at 2,000 ×g for 10 min and washed three times with 0.9% saline. Pellets were weighed to ensure exactly 40 mg/mL of CFT073 would be the stock solution for all studies described. CFT@ZIF was prepared by adding 1 mg of CFT073 from the stock solution and 500 uL of 1600 mM 2-methylimidazole (HMIM) into a 1.5 mL microcentrifuge tube, followed by the addition of 500 µL of 20 mM zinc acetate dihydrate (ZnOAC). The tube was capped, swirled for 20 s, and left to incubate at RT for 20 min. HMIM and ZnOAC solution were made using 100 mM NaCl solution to keep bacteria near isotonic conditions. The solution became cloudy after a few minutes and remained colloidal throughout the incubated timeframe. Longer time frames than 20 minutes led to free ZIF-8 as seen in Extended Data Fig. 2. After 20 min, the solution was centrifuged at 4500 *×g* for 10 min at 4°C. The supernatant was discarded, and the pellet was washed with ultrapure water twice. The final CFT@ZIF powder was either dried to characterize the sample or placed into 0.9% saline for *in vitro* and *in vivo* studies.

#### Bactericidal Assay

In a 1.5 mL microcentrifuge tube, either 0.9% saline or 500 mM acetic/acetate buffer pH 5 (Acetate buffer) at a volume of 975 µL was added to each tube. 25 µL of CFT073 or dried CFT@ZIF was added to each tube to obtain a 1 mg/mL concentration of bacteria. Acetate buffer can exfoliate the ZIF shell and was used as a control to show that untreated CFT073 will continue to grow when incubated for the same amount of time it takes to exfoliate CFT@ZIF. CFT@ZIF must be incubated for 30 min to be fully exfoliated. The untreated CFT073 or CFT@ZIF was suspended in 0.9% saline and left at 80 °C for 15 minutes. As previously stated, untreated CFT073 was exposed to identical stress conditions, dilutions, and incubation in 500 mM acetic/acetate buffer pH as CFT@ZIF biocomposites. Formalin-fixed CFT073 (CFT-Fixed) was made by placing 1 mg in 5% formalin to a final volume of 1 mL and incubating overnight. CFT-Fixed was then centrifuged at 4,000 ×g and washed with 0.9% saline, twice. Each of the tested conditions was serially diluted (10-2 to 10-9) and spotted on an LB agar plates to determine the colony-forming unit (CFU) titers formed after 12 h, individually. All experiments included n=3 per sample tested.

#### Agglutination Assay

Bacterial strains at an OD_600_ of 0.5 grown in Luria Bertani (LB) medium overnight at 37°C in static conditions. Saccharomyces cerevisiae (yeast) at an OD_600_ of 0.5 was grown in yeast extract-peptone-dextrose (YPD) medium overnight at 27 °C in an incubator at 225 rpm. Bacterial and yeast cultures were centrifuged at 2655 rcf for 5 minutes at room temperature and resuspended in 1000 µl of PBS..The OD of a 1 mg/mL stock of CFT073 was measured and used to adjust the concentration of the stock to a desired OD. 800 mM HMIM and 10 mM ZnOAC representing the final concentration in the CFT@ZIF reaction were prepared as samples. CFT-Heat was prepared by adding CFT073 to 100 mM saline for an OD 0.20 and final volume of 1 mL and submerged in an 80 °C water bath for 15 min. ZIF-8 was prepared in the same manner as CFT@ZIF but without the addition of CFT073. CFT@ZIF was prepared as mentioned above but CFT073 at an OD of 0.20 was added after HMIM and before ZnOAC. These were washed twice at 4,300 ×*g* and resuspended in 100 mM saline to 1 mL. CFT-Fixed was prepared as previously stated but also at an OD: 0.20. Each sample was added to yeast at a 2:1 ratio on a glass microscope slide and mixed by shaking for 2 min. Microscope images were taken at 10 × magnification at 10 min and 20 min.

#### Lactate dehydrogenase (LDH) Cytotoxicity Assay

Procedure was performed as recommended by vendor. Briefly, T24 and RAW 264.7 cells were seeded in a 96 well plate (100 uL/well) at a concentration of 1×10^6 cells/mL and incubated overnight in a 37 °C CO_2_ incubator. The next day the media was removed and replaced with clean media containing the designated CFT samples at the designated concentrations (100 uL/well) in triplicate and incubated for 4 h at 37 °C in a CO_2_ incubator. After 4 h 10 uL of lysis buffer was added to the negative control wells and incubated at 37 °C in a CO_2_ incubator for 30 mins. Next 100 uL of working solution was added to each well and incubated in the dark at room temperature for 30 mins. Lastly, 50 uL of stop solution was added to all wells and the plate was read at 490 nm.

#### Macrophage Uptake by Flow Cytometry

RAW Macrophage 264.7 cells were cultured in Dulbecco’s Modified Eagle Medium supplemented with 10% FBEssence and 1% penicillin-streptomycin (50 µg/mL). Cells were seeded at ∼10^5^ cells/mL in a 6 well plate one day prior to testing. The cells were incubated with 25 µg/mL of indicated samples for 4 hours. The cells were washed 3× with low pH buffer (0.5% acetic acid, 0.5 M NaCl, pH 3), 3× with 1× PBS, resuspended in 1 mL of 1× PBS, and transferred to a 5 mL sterile polystyrene tube. ∼10,000 gated events per sample were collected using a BD LSRFortessa™ flow cytometer. Raw data were processed and analyzed using FlowJo® software Version 10.6.1. Histogram overlays were normalized to mode to compare samples that varied in number of recorded events.

#### Macrophage Uptake by Fixed Cell Imaging

RAW Macrophage 264.7 cells were cultured in Dulbecco’s Modified Eagle Medium supplemented with 10% FBEssence and 1% penicillin-streptomycin (50 µg/mL). Cells were seeded at ∼10^5^ cells/mL in a 6 well plate one day prior to testing. The cells were incubated with 25 µg/mL of indicated samples for 4 hours. The cells were washed 3× with low pH buffer (0.5% acetic acid, 0.5 M NaCl, pH 3), 3× with 1× PBS, fixed with 4% paraformaldehyde, stained with 300 nM DAPI and 5 µg/mL WGA-TRITC, washed 3× with 1× PBS, washed 3× with autoclaved MilliQ water, and mounted on glass slides using Fluoroshield histology mounting media. Fixed cell imaging was performed with an Olympus FV3000 RS Confocal microscope. Raw images were processed using ImageJ software.

#### Antigen stimulation of splenic T lymphocytes

Mice were injected on days 0, 7, 14 and splenocytes were isolated from immunized mice 7 days after the third immunization. The cells were stained with anti-CD3-PacBlue, anti-CD4-PE/Cy7, anti-CD8a-FITC, and anti-B220-Alexa700 and analyzed by flow cytometry (gating strategy illustrated in **Fig S5A**). Another set of mice were vaccinated according to **Figure 2C**, at day 42 the spleens were collected and homogenized into single cell suspension. The cells were seeded at ∼ 1.0 × 10^6^ cells per well in a 24 well plate and supplemented with RPMI 1640 medium, 10 % FBEssence, 1 % penicillin-streptomycin, and 50µm β-mercaptoethanol. Cells were re-stimulated with 10 ug of untreated CFT073 (10 μg/mL) for 48 hours. The supernatant was tested for cytokine production by enzyme-linked immunosorbent assay (ELISA).

### Immunophenotyping of the draining lymph node

Mice were injected on days 0, 7, 14 and the draining lymph node was isolated from immunized mice 7 days after the third immunization. Briefly, the cells were homogenized into single cell suspension and stained at ∼ 1.0 × 10^6^ cells with anti-CD19-Alexa700, anti-CD95-APC, anti-GL7-Alexa488, anti-CD86-BV785, anti-CD184-BV421, and anti-IgG1+-PE and analyzed by flow cytometry (gating strategy illustrated in **Fig. S5B**). B cells in the light zone express CD86^hi^ and B cells in the dark zone express CD86^low^.

### Antibody and cytokine ELISA

CFT073-specific antibody production was determined by following a previously published method.^81^ Briefly, lyophilized CFT073 was resuspended in 0.05 M pH 9.6 sodium carbonate/bicarbonate buffer to a concentration of 0.31 mg/mL, 150 µL was added to each well, and was incubated at 37°C for 90 min. The plate was emptied and washed 4 times with wash buffer. 200 µL of blocking buffer was added to each well and was incubated at 37°C for 45 min. The plate was emptied and washed 4 times with wash buffer. Mouse serum was serially diluted 7 times starting at a 200× dilution using 1x PBS at ph 7.4, 150 µL added to each well, and incubated at 37°C for 90 min. The plate was emptied and washed 4 times with wash buffer. Alkaline phosphatase-conjugated goat anti-mouse IgG in conjugate buffer was added 150 µL per well and incubated at 37°C for 90 min. The plate was emptied and washed 4 times with wash buffer. 1 mg/mL p-nitrophenylphosphate in substrate buffer was added at 150 µL per well and the plate developed for 15 min at R.T. The plate was read at 405 nm and the absorbance values of the buffer blank wells averaged and subtracted from the entire plate. The blank-subtracted values of each mouse group were reported as the average ± standard deviation for each dilution. The levels of TNF-α, IL-17, IL-4, IL-2, and IFN-_ were determined by ELISA following protocols recommended by the manufacturer.

### Quantitation of Immunoglobulin Isotyping

Assay was performed following guidelines provided by the vendor. Further, assay was used to quantify 6 immunoglobulins from frozen serum samples. Briefly, 2µL serum was diluted to 50,000 times using assay buffer and IgG1, IgG2a, IgG2b, IgG3, IgA and IgM levels were measured according to the manufacturer’s protocol. Streptavidin phycoerythrin (SA-PE) intensity was analyzed by a FACS LSR Fortessa instrument (Becton Dickinson), and each analyte was quantified relative to the kit standard curve using LEGENDplex software, version 8.0.

### Animal Studies

#### Ethics Statement

Female BALB/c were obtained from Charles River Lab (Wilmington, Ma). All animal studies were done in accordance with protocol #19-06 approved by the University of Texas at Dallas Institutional Animal Care and Use Committee (IACUC).

#### Vaccinations

20 BALB/c mice were divided into five groups (n = 4) and injected with saline, formalin CFT-Fixed, CFT@ZIF, CFT-Heat, or CFT@ZIF-Heat suspended in saline. Heat-treated samples were placed in 80°C water for 15 minutes. CFT073 solutions were prepared such that 100 µL delivered 10 µg of CFT073. Doses of 100 µL of saline, CFT-Fixed, CFT@ZIF, CFT-Heat or CFT@ZIF-Heat were administered subcutaneously on day 0, 7, and 14, and blood was withdrawn submandibularly on day 0, 14, 21, and 42. The blood was centrifuged to remove cells, and the anti-CFT073 IgG content of the resultant serum was determined by ELISA as described above. At the end of the study, the mice were sacrificed for histological analysis on the spleen, liver, kidney, lung, heart, and the skin at the administration site. The mice were sacrificed by carbon dioxide asphyxiation, the organs collected, and fixed in 4% formaldehyde overnight. The fixed organs were moved to a 70% ethanol solution and processed with an ASP300 S tissue processor (Leica Biosystems, Buffalo Grove, IL) for dehydration into paraffin. The organs were then embedded into paraffin wax using a HistoCore Arcadia C and H paraffin embedding station (Leica Biosystems, Buffalo Grove, IL). Each organ was sliced into 4 µm sheets using a RM2235 manual microtome (Leica Biosystems, Buffalo Grove, IL) and imaged with a DMi1 optical microscope (Leica Biosystems, Buffalo Grove, IL) at 40×magnification.

#### Body Clearance

10 BALB/c mice were fed a non-fluorescent diet and shaved 12 hours before imaging to reduce autofluorescence from the hair. The mice were anesthetized with isoflurane and injected with 100 µL saline (n=4), CFT(smURFP)-Fixed, or CFT(smURFP)@ZIF. The CFT073 containing solutions were prepared such that 100 µL delivered 10 µg of CFT073. A series of time points were taken after injection at 30 min, 2 h, 4 h, 8 h, 18 h, 24 h, 48 h, 72 h, 120 h, 168 h, 216 h, and 288 h. The fluorescence returned to the levels of saline at 48 hours for CFT-Fixed and 216 h for CFT@ZIF.

#### CFT073 Sepsis Challenge

As ours is a developing technology with many unknowns, we expect that for the CFT@ZIF there will initially be larger variances in humoral response which we expect to be around 20%. To detect at least a 20% difference in the protein expression changes with a power = 0.8 and a significance level of 0.05, a sample size of at least 6 mice will be necessary. 20 BALB/c mice were divided into five groups (Two cohorts, n=3 and n=4) and injected with saline, formalin CFT-Fixed, CFT@ZIF, CFT-Heat, or CFT@ZIF-Heat suspended in saline. Heat treated samples were placed in 80°C water for 15 minutes. CFT073 solutions were prepared such that 100 µL of 0.9% saline delivered 10 µg of CFT073. Doses of 100 µL of saline, CFT-Fixed, CFT@ZIF, CFT-Heat or CFT@ZIF-Heat were administered subcutaneously on day 0, 7, and 14. On day 21, all mice were injected interperitoneally with a lethal dose of 3.6 × 10^8^ CFU of CFT073 per mouse and monitored for 48 h. All mice were euthanized when they became moribund, which is defined by lack of movement for over 15 minutes and when gently touched and shaking in place. After a mouse was euthanized, the spleen and liver were collected, homogenized, and plated onto LB agar plates at dilutions ranging from 10^−2^ – 10^−9^ to determine the CFU/g. The blood was also taken from each mouse and serially diluted in PBS to determine the CFU/mL.

## Acknowledgements

This project was partially funded by The University of Texas at Dallas Office of Research through the SPIRe grant program. J.J.G. thanks the National Science Foundation [CAREER DMR-1654405 and DMR-2003534] and the Welch Foundation [AT-1989-20190330]. N.J.D. thanks the Welch Foundation [AT-2030-20200401]. The Gassensmith Lab would like to thank the University of Texas at Dallas lab animal resource center (LARC) for mouse maintenance. We especially like to thank Tyler Tornblom and Bradly Woody from LARC for the mice training they provided. C.E.B. thanks the National Science Foundation Graduate Research Fellows Program (1746053).

## Author Contribution

Primary manuscript writing and editing was done by M.A.L., M.A.I. N.J.D. and J.J.G. Cytokine ELISA, body clearance, flow cytometry, and survival challenge were done by M.A.L. SEM, EDX, Mouse injections and blood draws were done by M.A.L. and F.C.H. TEM imaging was done by O.R.B. Confocal images were taken by O.R.B. and C.E.B. Cytotoxicity assays were done by O.R.B and R.E. PXRD was done by Y.W. smURFP induction was done by K.V. Histological analysis was done by A.S, F.C.H, and J.G. Whole IgG ELISA was done by S.P and Isotype multiplex assay was done by J.G. Flow prep was done by M.A.L., A.S., and O.R.B. ZIF optimization was done by M.A.L. and F.C.H. Stabilization studies over time were done by F. C. H. Agglutination assay was conducted by T.H. and S.V. Bacteria growth and killing assay was done by F.C.H. Immunological studies were aided by M.D.B. Funding was raised by N.J.D. and J.J.G.

