## Supporting Information for "Metal-Organic Framework Encapsulated Whole-Cell Vaccines Enhance Humoral Immunity against Bacterial Infection"

A Whole Cell Metal-Organic Framework Encapsulated Vaccine Against Septicemic UPEC Infection.

**Supplemental Data**

**Biolegend ELISA Kits**

IFN-γ: Sensitivity 4 pg/mL and Standard range 15.6 – 1,000 pg/mL

TNF-α: Sensitivity 4 pg/mL and Standard range 7.8 – 500 pg/mL

IL-17: Sensitivity 8 pg/mL and Standard range 15.6 – 1,000 pg/mL

IL-4: Sensitivity 1 pg/mL and Standard range 2.0 – 125 pg/mL

IL-2: Sensitivity 1 pg/mL and Standard range 2.0 – 125 pg/mL

IL-6: Sensitivity 2 pg/mL and Standard range 7.8 – 500 pg/mL

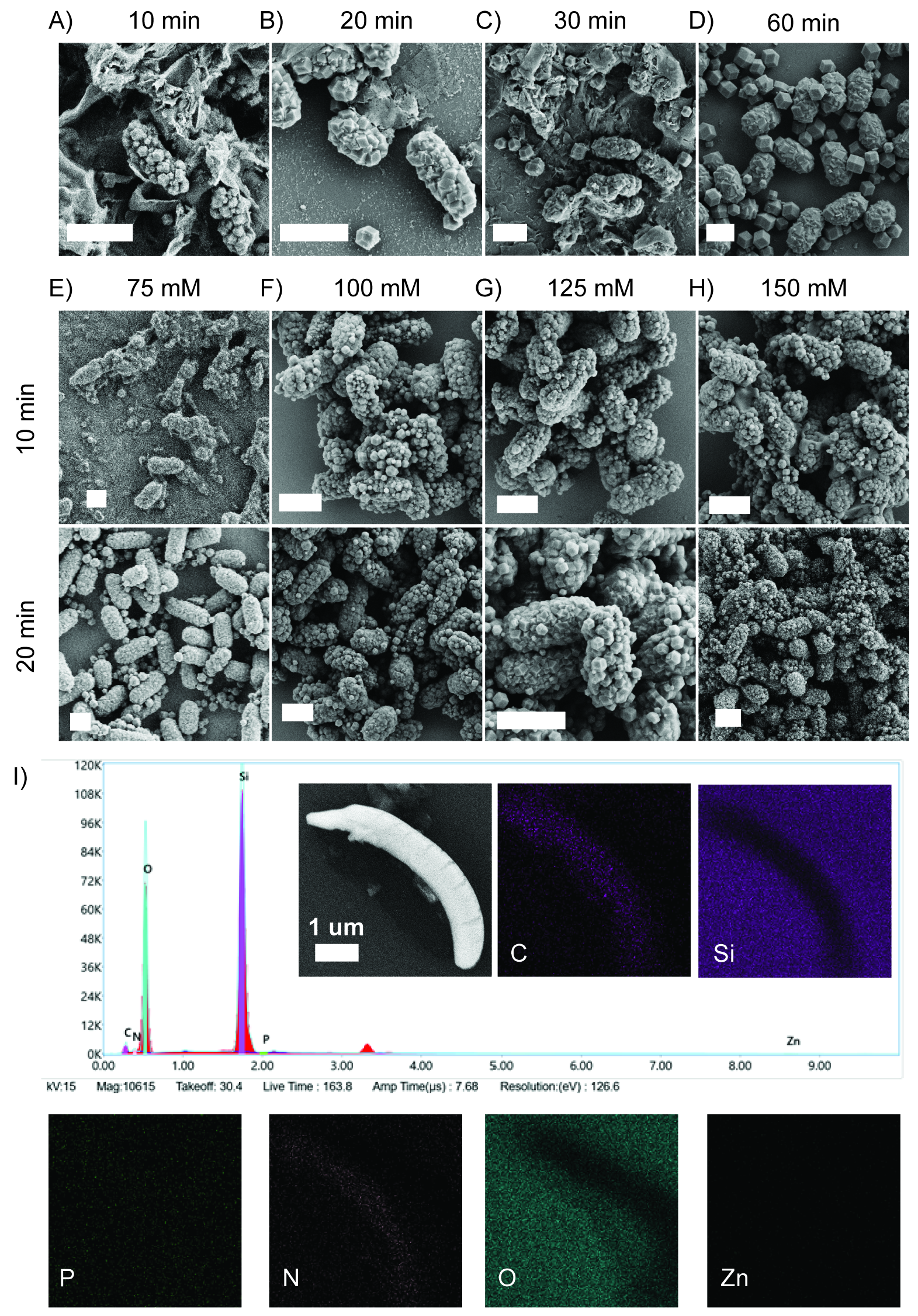

**Fig. S1 Shorter incubation times and saline reduce the amount of free ZIF-8.** Live CFT073 was incubated with an aqueous solution of zinc and 2-methylimidazole and left at room temperature for A) 10 minutes, B) 20 minutes, C) 30 minutes, or D) 60 minutes. After each time point, the sample was centrifuged to remove excess zinc and 2-methylimidazole and washed with water three times. CFT@ZIF encapsulation was performed with 1 mg of CFT073 in E) 75 mM, F) 100 mM, G) 125 mM, or H) 150 mM saline solutions for either 10 or 20 minutes to reduce cell bursting and the amount of free ZIF-8. I) Element distribution of CFT073 by EDX. The graph shows the presence of carbon, oxygen, nitrogen, and phosphorus. Image maps show carbon and nitrogen signals come from CFT073. White scale bars are 2 µm, unless stated otherwise.

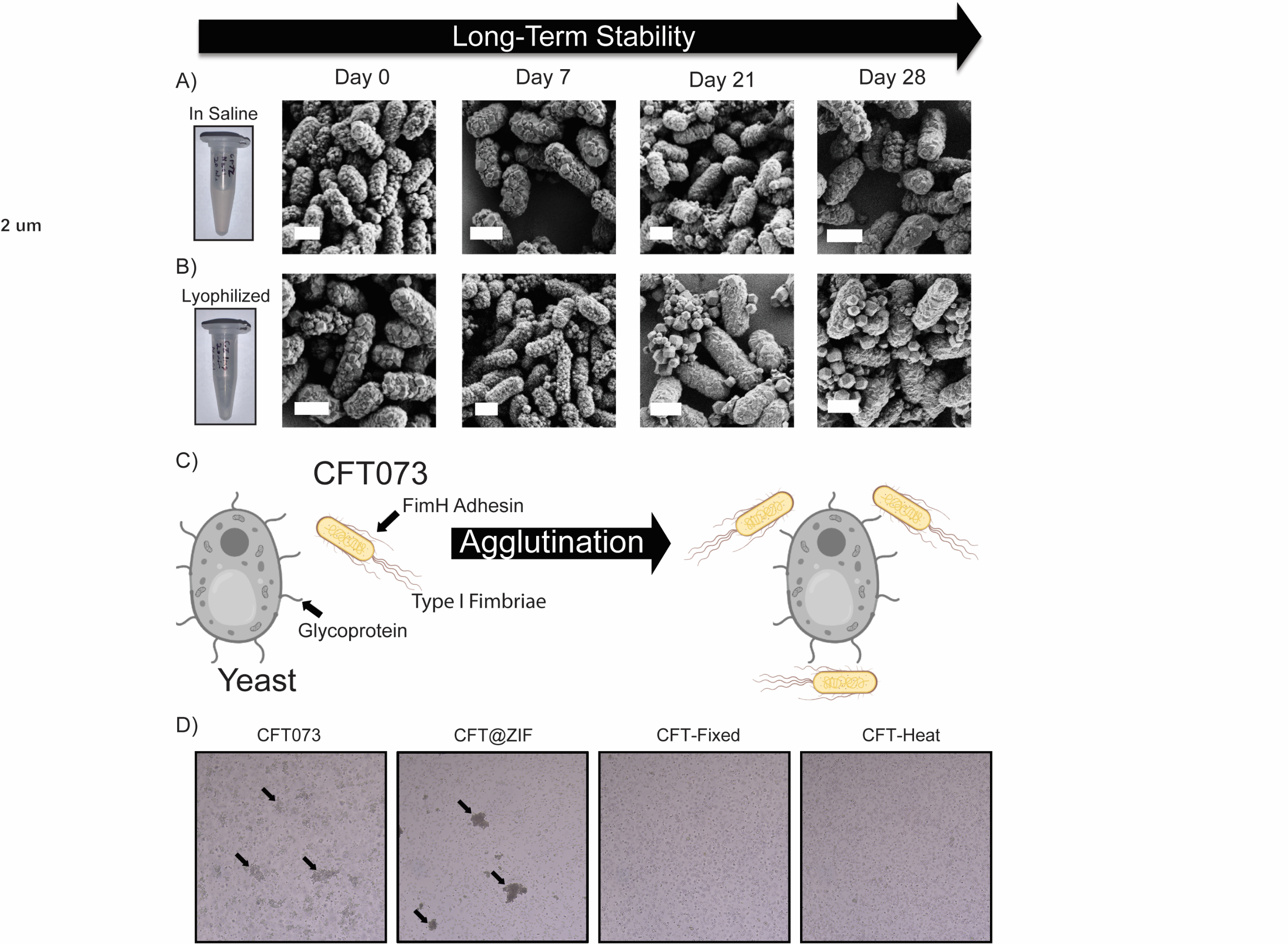

**Fig. S2 CFT@ZIF Stability and Agglutination Assay.** The stability of CFT@ZIF in A) 0.9% saline solution and B) after lyophilization. Both formulations were imaged by SEM at Day 0, 7, 21, and 28. White scale bars are 2 µm. C) An illustration of the principle of agglutination assay. In this qualitative assay the FimH adhesin protein in type 1 fimbriae of CFT073 bind sugar moieties, specifically mannose. *Saccharomyces cerevisiae* yeast contain large amounts of mannose in their outer cell wall in the form of glycoprotein mannoprotein. This interaction forms large, crosslinked clumps called agglutinates which are clearly visible in light microscopy. D) The agglutination assay was tested at 1.0 × 10^8^ CFU/mL (OD = 0.2) for CFT073 and CFT@ZIF and 2.0 × 10^8^ CFU/mL (OD = 0.4) for CFT-Fixed and CFT-Heat and shows only CFT073 and CFT@ZIF retains binding with yeast. Whereas, CFT-Fixed and CFT-Heat lose binding after treatment.

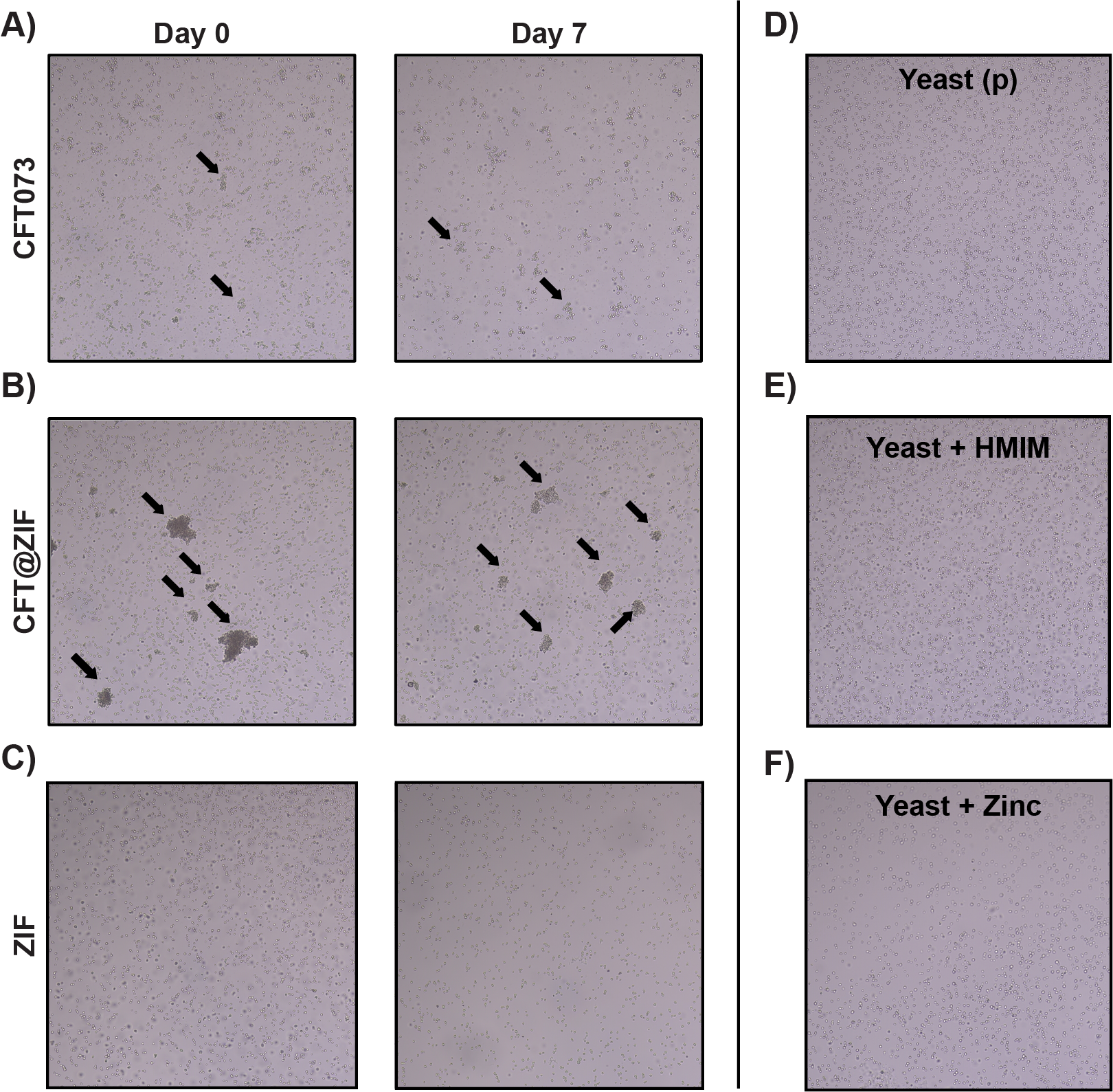

**Fig. S3 Testing the Agglutination at Different Time Points.** Agglutination assays were conducted to determine the stability of our formulation over the course of 7 days. We kept A) CFT073, B) CFT@ZIF, and C) ZIF in 100 mM saline and performed an agglutination assay at day 0 and kept the test samples at RT without shaking for 7 days, then repeated the agglutination assay on the 7-day old samples. D) Images of pristine yeast in saline, E) yeast incubated with 800 mM HMIM, and F) yeast incubated with 10 mM ZnOAC to show the individual precursors of ZIF don’t cause agglutination.

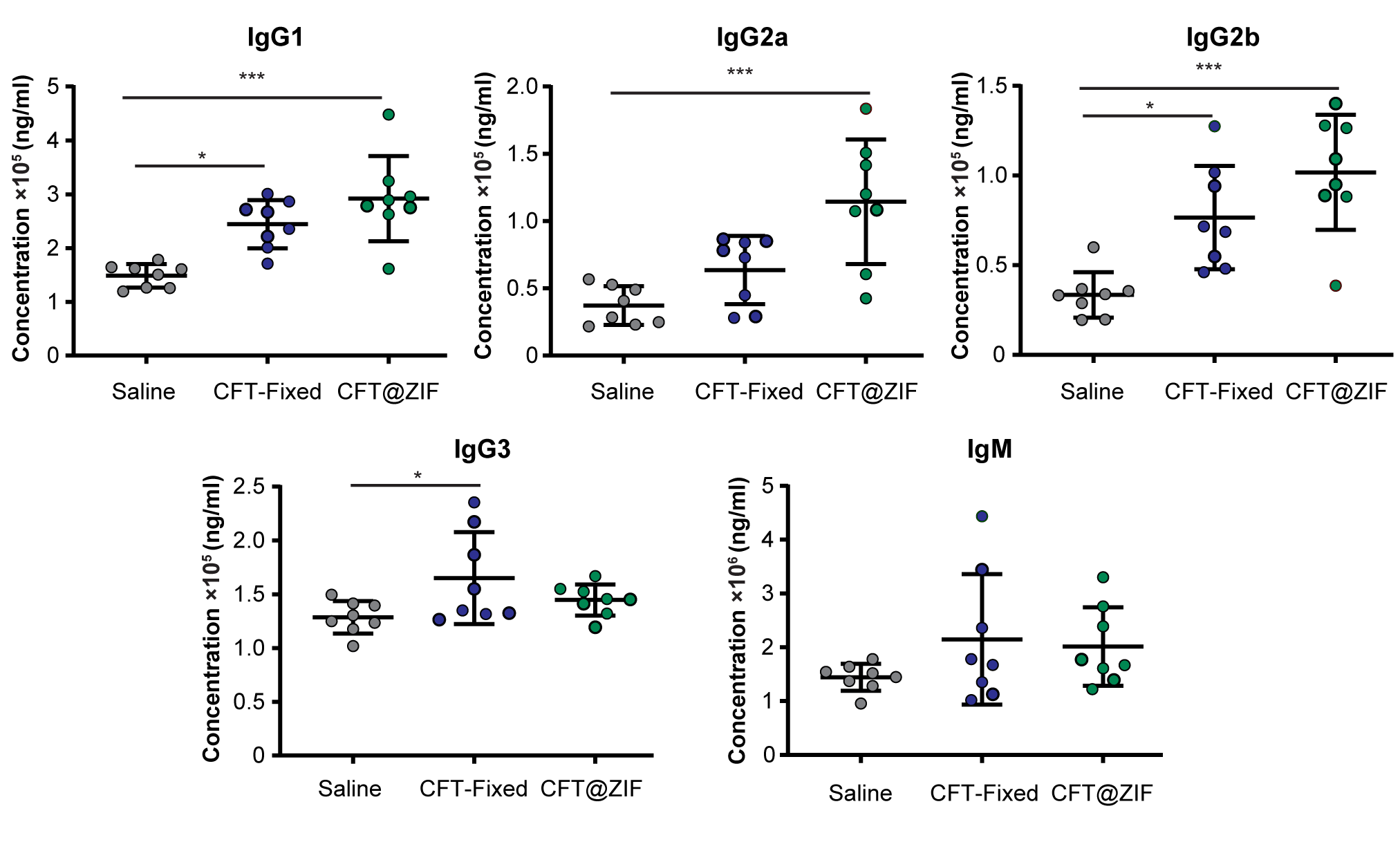
**Fig. S4 IgG Subclass Production.** Mice (n=8) were vaccinated following the schedule in **Figure 2C** and on day 21 blood was collected to obtain serum. The antibody subclasses IgG1, IgG2a, IgG2b, IgG3, and IgM was measured. Statistical signiﬁcance was calculated using two-way ANOVA with Tukey’s multiple comparison post-test (*p < 0.05, **p < 0.01, ***p < 0.0005, ****p < 0.0001).

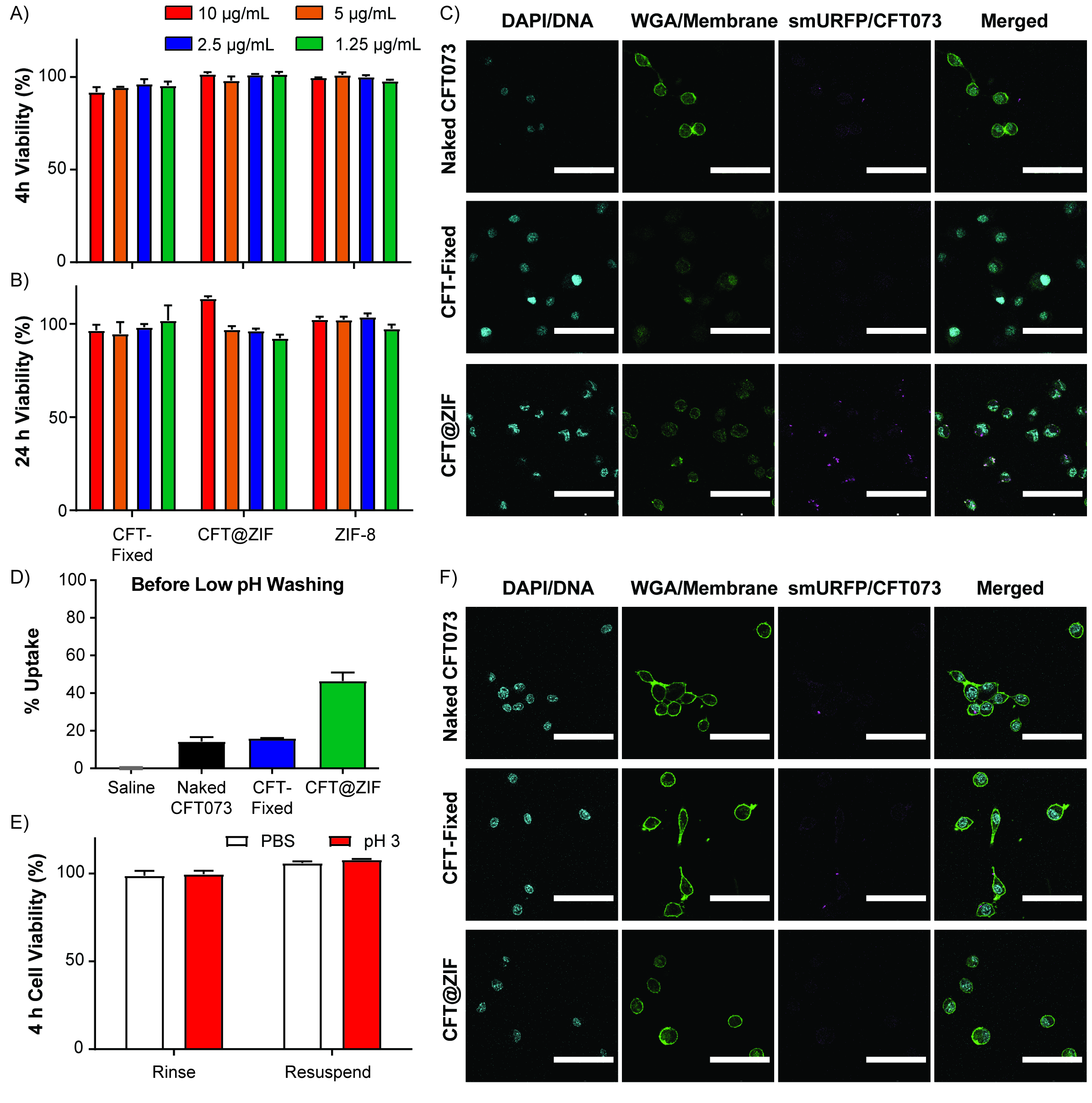

**Fig. S5 Viability and CFT@ZIF low pH wash study.** The viability of CFT-Fixed, CFT@ZIF, and ZIF was tested at different concentrations on RAW 264.7 macrophages for A) 4 h and B) 24 h. Before low pH wash C) Representative confocal micrographs (N=10) of smURFP expressing naked CFT073, CFT-Fixed, and CFT@ZIF incubated with RAW 264.7 macrophages for 4 h. Scale bar = 50 µm. D) Bar graphs of flow cytometry smURFP fluorescence (N=3) without pH 3 washing of cells measure both surface bound and internalized bacteria. E) We tested the pH 3 washing on the RAW 264.7 macrophages to confirm the washes would not be toxic to the cells. The rinse group was just incubated with pH 3 solution for 5 min and the resuspension group used a 1 mL pipettor to aspirate the pH 3 solution 10 times, followed by a 5 min incubation. Both samples were centrifuged to remove pH 3 solution and another aliquot of pH 3 solution was added—this was repeated for a total of 3 times. The next set of data is following washing the cells at low pH to remove surface-bound bacteria: F) Representative confocal micrographs (N=10) of smURFP expressing naked CFT073, CFT-Fixed, and CFT@ZIF incubated with RAW 264.7 macrophages for 4 h and washed with pH 3 buffer. Scale bar = 50 µm

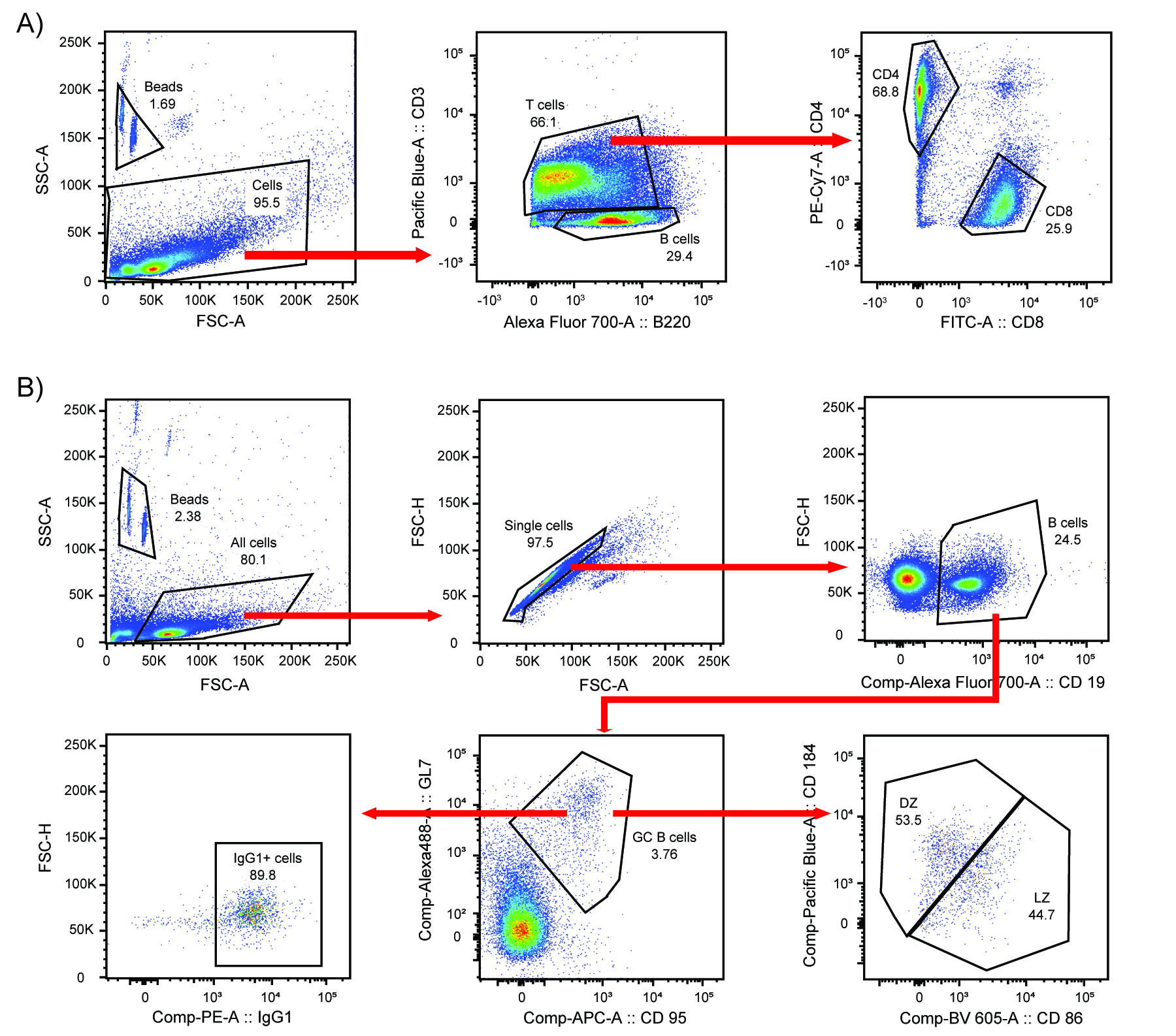

**Fig. S6 Representative gating strategy.** A) For adaptive immune cells: T cells were defined as CD3+ cells. Helper T cells were defined as CD3+ CD4+, Cytotoxic T cells were defined as CD3+ CD8+, and B cells were defined as B220+. B) For germinal center: B cells were defined as CD19+ and GCBCs were defined as CD19+ CD95+ GL7+.
